# Deletion of NLRP3 gene blocks traumatic brain injury induced abnormal immune response in 3xTg AD mice

**DOI:** 10.64898/2026.02.11.705384

**Authors:** Jakob Green, Zheng Liu, Sarah Timis, Christopher Nelson, Alexandra Pedin, Chunqing Guo, Xiang-Yang Wang, Shijun Zhang, Dong Sun

**Author notes:** **Corresponding author:** Dong Sun, M.D, PhD.

## Abstract

Traumatic brain injury (TBI) is a significant risk factor for the development of Alzheimer’s disease (AD) and related dementia. In both TBI and AD, inflammation plays a pivotal role. It is known that NLRP3 inflammasome plays an important role in pathogenesis of AD while TBI triggers activation of NLRP3 inflammasome in the brain. To evaluate the importance of NLRP3 inflammasome in mediating TBI-induced immune response in the context of predisposition of AD, we have examined the immune profiles in the brain using a novel transgenic AD mouse line with the NLRP3 gene deleted. Briefly, a group of 4 months old male and female 3xTg and 3xTg/NLRP3^-/-^ mice received a moderate lateral fluid percussive injury or sham. Immune cell phenotypes and cytokine gene expression were assessed at 3- and 7-days post injury (dpi). We found that NLRP3 gene deletion counteracted injury-induced alteration of immune response in AD mice with a significant sex-related difference. Specifically, TBI induced a significant brain infiltration of neutrophils, macrophages and γδ T-cells in 3xTg mice in both sexes at 3dpi, and NLRP3 gene deletion blocked this injury effect only in males not in females. NLRP3 gene deletion also blocked injury-enhanced IL-1β, TNF-α gene expression in male AD mice, but not in female mice. In conclusion, our study has confirmed that TBI significantly alters the immune response in 3xTg-AD mice and NLRP3 inflammasome is important in mediating these TBI-induced changes with significant sex-related differences.

## Introduction

The intersection of traumatic brain injury (TBI) and Alzheimer’s disease (AD) is a topic of much discussion due to the increased risk of developing AD after TBI (Julien et al., 2017). TBI is the leading cause of death and disability for young people worldwide (Faul and Coronado, 2015). Following TBI, the primary injury induces irreversible and untreatable brain tissue damage. The subsequent secondary injury causes progressive neurodegeneration and delayed cell death (Loane and Faden, 2010; McIntosh et al., 1996). Extensive studies have established that TBI leads to cognitive decline and dementia, and growing body of evidences have suggested that TBI accelerates AD development (Gardner et al., 2014; Gardner and Yaffe, 2014; Iacono et al., 2021; LoBue et al., 2017; Schaffert et al., 2018). According to the United States Centers for Disease Control and Prevention, individuals with a history of moderate TBI have a 2-3 times higher risk of developing AD, and studies reported that TBI can lead to an earlier onset of AD pathology in as much as 4 to10 years earlier (Gardner et al., 2014; Gardner and Yaffe, 2014; Iacono et al., 2021; LoBue et al., 2017; Schaffert et al., 2018). Furthermore, studies have also shown that TBI accelerates beta-amyloid (Aβ) deposition and tau abnormalities, two pathological hallmarks of AD, and cognitive impairment (Johnson et al., 2012; Roberts et al., 1994).

Although those extensive epidemiological evidences have suggested that TBI accelerates the onset and severity of AD, the underlying mechanisms linking TBI to AD remain unclear. Among possible factors, the inflammatory response is largely recognized as the major culprit as it is a common and essential player in both TBI and AD, and TBI-induced neuroinflammation persists for many years (Hellewell et al., 2016; Krstic and Knuesel, 2013; Kumar and Loane, 2012; Lozano et al., 2015; Pimplikar, 2014). Following TBI, there is a robust inflammatory response characterized by infiltration of immune cells, activation of resident glial cells, and an increase in the levels of inflammatory mediators including cytokines and chemokines, such as IL-1β, IL-18 and TNF-α (Loane and Byrnes, 2010; Morganti-Kossmann et al., 2007; Smith et al., 2013; Ziebell and Morganti-Kossmann, 2010). TBI-induced inflammatory responses persist long after the initial injury, up to 47 years as reported by Smith et al (Smith et al., 2013). Accumulating evidence also suggests the immune/inflammatory response as an essential driver for AD development. Pathologically, chronic glial activation and elevated levels of proinflammatory cytokines were observed in AD models and patients (Heneka et al., 2015; Morimoto et al., 2011; Parachikova et al., 2007; Prokop et al., 2013). Such chronic inflammation can induce dysfunction of microglia causing inefficient clearance of Aβ species that is supported by the reported strong correlation between chronic inflammation, Aβ deposition, and tau hyperphosphorylation (Lee et al., 2010; Mawuenyega et al., 2010; Rojo et al., 2008; Sy et al., 2011).

It is known that innate immunity plays a major role in inflammatory responses following TBI (McKee and Lukens, 2016; Nizamutdinov and Shapiro, 2017). Inflammasomes are critical regulators of the innate immune response to pathogen-associated and danger-associated molecular patterns (Schroder et al., 2010). Recently the NOD-like receptor family pyrin domain containing 3 (NLRP3) inflammasome has been identified as a critical multiprotein platform regulating the innate immune response and the production of pro-inflammatory cytokines (Martinon et al., 2002). Notably, emerging evidences suggest that TBI induces activation of the NLRP3 inflammasome and is involved in the pathological development of AD (Couturier et al., 2016; Freeman and Ting, 2016; Halle et al., 2008; Heneka, 2017; Heneka et al., 2013; Irrera et al., 2017; Irrera et al., 2020; Latz et al., 2013; Liu et al., 2013; Ma et al., 2017; Saresella et al., 2016; Tan et al., 2013; Wallisch et al., 2017; Walsh et al., 2014). Activation of NLRP3 is induced by multiple stimuli, including reactive oxygen species (ROS), mitochondrial damage, ATP, and potassium ion efflux from injured cells following tissue damage such as TBI (Latz et al., 2013; Mariathasan et al., 2006; Pelegrin and Surprenant, 2007). Following TBI, increased activation of the NLRP3 inflammasome in the injured brain is observed in TBI models (Kuwar et al., 2019; Lin et al., 2017; Liu et al., 2013; Xu et al., 2018), and inflammasome-induced cell death contributes to secondary tissue damage (Adamczak et al., 2014). This is supported by clinical evidence that proteins in the NLRP3 inflammasome complex and downstream effectors including IL-1β and IL-18 are found in the CSF of TBI patients and the increased levels of which are associated with poor prognosis (Adamczak et al., 2012; Wallisch et al., 2017). Animal studies have also found that NLRP3^-/-^ mice have preserved cognition with less brain damage and inflammation following TBI (Irrera et al., 2017).

Studies have also supported the importance of the NLRP3 inflammasome in AD. In both AD animal models and patients, the protein expression levels of NLRP3, ASC, caspase-1 and IL-1β are upregulated (Couturier et al., 2016; Halle et al., 2008; Heneka et al., 2013; Saresella et al., 2016; Tan et al., 2013). Both IL-1β and IL-18 have shown essential roles in AD pathological development including synaptic plasticity, amyloidogenesis, and tauopathy (Cumiskey et al., 2007; Ghosh et al., 2013; Griffin et al., 1989; Lynch, 2015; Nicoll et al., 2000; Zhu et al., 1999). NLRP3^-/-^ and Caspase-1^-/-^ mice carrying mutations associated with familial AD (APP/PS1) have exhibited reduced Aβ burden, preserved synaptic integrity, and improved spatial memory functions (Heneka et al., 2013). Studies in 5xFAD mice carrying the ASC^+/-^ genotype also support this notion (Couturier et al., 2016). Conversely, NLRP3 over-activation could promote differentiation of T cells into pro-inflammatory Th1 and Th17 effector phenotypes contributing to sustained inflammation (Gris et al., 2010), while deletion of NLRP3 in aged mice provides protection from aging related cognitive decline (Youm et al., 2013). Furthermore, studies have revealed that Aβ plaques induce activation of NLRP3 inflammasome, which in turn further exacerbates neuroinflammation and drives tau pathology (Halle et al., 2008; Ising et al., 2019).

Based on these findings, we speculate that TBI-induced abnormal activation of the NLRP3 inflammasome and downstream chronic inflammation acts as a mechanistic crosslink between TBI and AD accelerating the onset and/or severity of AD in patients with TBI history. In this study, utilizing a novel transgenic mouse model with the NLRP3 gene deleted at the base of a widely used triple transgenic AD mouse line (3xTg), we have investigated the importance of the NLRP3 inflammasome in regulating post-TBI immune/inflammatory response in the context of AD predisposition at the acute and subacute stages following TBI in both male and female animals. We have found that TBI altered the immune response in 3xTg mice and NLRP3 gene deletion counteracted the injury effect with a significant sex-related difference.

## Materials and Methods

### Experimental Animals

The triple transgenic AD (3xTg) mouse line carrying the APP695 gene with Swedish mutations (KM670/671NL/M596L), PSEN1 mutation (M146V), and tau mutation (P301L) (Jax. Stock No: 004807) were used. To determine the importance of NLRP3 inflammasome in mediating post-TBI AD development, we have created a unique transgenic AD mouse line with NLRP3 gene knock out (homozygous 3xTg with NLRP3 gene null). This line was derived by crossing homozygous 3xTg mice with NLRP3^-/-^ mice (Jax. Strain No: 021302). Double transgenic offspring were then crossed to establish the breeder line. The VCU Transgenic Core performed breeding and genotyping of the 3xTg, 3xTg/NLRP3 null mouse lines. Animals were housed in a vivarium with controlled temperature, humidity, and were kept on a 12-hour light-dark cycle with food and water provided *ad libitum*. All procedures were approved by the Institutional Animal Care and Use Committee.

### Surgical Procedure

At the age approximately 4 months old, animals were randomly divided into groups receiving either a moderate lateral fluid percussion injury (LFPI) or sham surgery. For female animals, estrogen cycle was monitored by visual observation following the published method (Byers et al., 2012) to ensure animals were injured at the same estrogen cycle at diestrus stage. The LFPI produces a mixed focal and diffuse axonal injury. The surgical instruments were sterilized and aseptic procedures were followed. Briefly, the mouse was first anesthetized in a plexiglass chamber with 4% isoflurane, then fixed to a stereotaxic frame and ventilated with 2.5% isoflurane mixed with 30% oxygen for the duration of the surgery. A continuously heated water pad was placed under the mouse during surgery to protect from hypothermia. After a midline incision on a shaved head to expose the skull, a craniotomy was made to the left of the middle line suture halfway between lambda and bregma using a 2.7mm trephine. A Luer-lock hub made with a 20-gauge needle cap was affixed to the craniotomy site with cyanoacrylate and dental acrylic. Anesthesia was turned off and the animal was allowed to recover from surgery in a warm cage for 2 hours before being subjected to TBI. For injury, the mouse was re-anesthetized with 4% isoflurane, hub filled with 0.9% saline and attached to a recalibrated FPI instrument. When the anesthesia wore off, the mouse was injured at the moderate injury level with the fluid pulse at 1.80±0.05 ATM. Immediately after injury, the mouse was placed on a heated pad and righting time was recorded. After righted, the mouse was re-anesthetized, the Luer-lock syringe hub was removed, and skin incision was sutured. The animal was returned to a clean cage on a warm blanket until fully recovered before being returned to the animal care facility. Sham group animals went through the same surgical procedure without receiving the fluid pulse. All animals received post-operative care.

To examine the effect of TBI on the immune cell response, a total of 43 3xTg mice (male: n= 20; female: n=23) and 41 3xTgNLRP3^-/-^ mice (male: n=20; female: n=21) were included in the final data collection (10% attrition rate in our TBI animals). Animals were sacrificed at 3- or 7-days post-injury (3 or 7 dpi). Mice were deeply anesthetized with an overdose of isoflurane inhalation followed by a transcardial perfusion with 30mL of ice-cold PBS. The brains were then quickly dissected on ice. After the whole brains were extracted, a small piece (1×1 mm^2^) of cerebral cortex close to the injury site was dissected for quantitative PCR, the rest of brain was dissociated for flow cytometric analyses (FACs). To confirm the efficacy of NLRP3 gene deletion, 6 male 3xTg mice (n=3/sham/TBI) and 8 male 3xTgNLRP3^-/-^ mice (n=4/sham/TBI) were perfused with ice-cold PBS at 3 dpi, cerebral cortex dissected and homogenized for Western blotting analysis.

### Flow cytometry

The frequencies of myeloid cell populations and T lymphocyte subsets were examined using multi-color FACs as we published previously (Yu et al., 2021). Briefly, the dissected individual brain was digested using the Neural Dissociation Kit (P) from Miltenyi Biotec to prepare single cell suspensions following the manufacturer’s guidelines. The obtained single cells were washed in phosphate-buffered saline (PBS) and resuspended in 40% Percoll (Sigma-Aldrich). The cell suspension was gently overlaid onto 70% Percoll and centrifuged for 20 min at 400×g to enrich mononuclear cells (MNCs). MNCs from the brain were incubated in phorbol 12’-myristate-13’-acetate (PMA, 10 nM, Sigma Aldrich) plus ionomycin (0.5 µM, Sigma Aldrich) for 3 hours, followed by treatment with brefeldin A (BFA, 5 µg/mL, Biolegend) to block cytokine secretion for another 2 hours. Cells were then surface stained with a panel of surface antibodies including CD11b (M1/70, #101236), CD45 (30-F11, #103147), Ly6G (1A8, #127606), CD4 (GK1.5, #100412), gdTCR (UC7-13D5, #107508), after fixing on ice for 30 min, cells were permeabilized and stained with IL-17A (TC11-18H10.1, #506920). All antibodies were purchased from Biolegend (San Diego, CA). The cells were analyzed using LSRFortessa™ Cell Analyzer (BD Biosciences).

### Real-time PCR

Real-time quantitative PCR (RT-qPCR) analysis of inflammatory cytokines was performed as we previously described (Yu et al., 2021). Briefly, total RNA was extracted from brains using TRIzol Reagent (ThermoFisher Scientific, Whaltham, MA). Reverse transcription and quantitative PCR were conducted using carboxyfluorescein (FAM)-labeled Taqman probe sets from ThermoFisher Scientific. Gene transcription of *il1b* (#Mm01336189_m1), *tnfa* (#Mm00443258), *IL-18* (#Mm00434225_m1), and *IL-17f* (Mm00521423_m1) was quantified relative to the expression of β-actin (#432933E), and normalized to that measured in control groups by standard 2(-ΔΔCt) calculation.

### Western blotting

For protein analysis, the ipsilalteral cortex was homogenized with RIPA buffer (Stock 10x RIPA, 20-188, EMD Millipore, MA), 0.1% SDS (Bio-Rad, 1610416), 1% Triton X-100 (Sigma-Aldrich, T8787), mini complete cocktail (Roche, 11836170001), 1mM EDTA (Quality Biological, 351-027-721). Homogenates were centrifuged at 16,000g at 4°C for 25 minutes and supernatants were collected and stored at -80°C until use. The total protein concentration was determined using the BCA method (Pierce, Rockford, IL). Amount of 20µg protein for each sample was boiled in Laemmli buffer (Bio-Rad, 1610747) and resolved using SDS-PAGE on a 4-20% Criterion TGX gel (Bio-Rad, 5671091). After SDS-PAGE, protein was transferred to a low-fluorescence PVDF (Bio-Rad) and blocked with either 5% BSA in tris-buffered saline with 0.1% Tween 20 (TBST) or 5% milk in TBST. The following primary and secondary antibodies were used: anti-NLRP3 (1:800, Cell Signaling Techonology, 15101S), anti-rabbit HRP (1:1000, Santa Cruz Biotechnology, sc-2357). The membranes were then washes and imaged using ChemiDoc MP imaging system (Bio-Rad, USA). The analysis of the images was done using the Image Lab 6.0 software (Bio-Rad, USA).

### Statistical Analysis

All the data were expressed as means ± SEM when applicable. The differences among multiple groups were assessed using a 2-way analysis of variance (ANOVA). A Tukey’s post hoc comparison between groups was used to determine specific group significance. Probability values less than 0.05 (p<0.05) were considered significant. Statistics were performed using JMP Pro 16 (JMP statistical software) and images prepared in GraphPad Prism 9.

## Results

### TBI alters the innate immune response in 3xTg mice and loss of NLRP3 counteracts the injury effect in a sex dependent manner

We have previously reported that TBI enhances expression of NLRP3 and downstream signaling elements at the acute and subacute stages following a focal brain injury in rats (Kuwar et al., 2019). In this study using Western blotting, we first assessed the expression level of NLRP3 in cerebral cortex in a group of male 3xTg and 3xTg/NLRP3^-/-^ mice to confirm the injury effect and the efficacy of NLRP3 gene deletion. At 3 days post injury (dpi), injured 3xTg mice had significantly higher expression of NLRP3 than the matched sham control, whereas NLRP3 was absent in 3xTg/NLRP3^-/-^ mice in both sham and TBI groups (**Supplemental Fig. 1**). This confirmed that TBI-enhanced NLRP3 expression in 3xTg mice was abolished in 3xTg/NLRP3^-/-^ mice.

To evaluate the influence of TBI on the immune response and the importance of NLRP3 in mediating this process, we compared the immune cell profiles in the brains of 3xTg and 3xTg/NLRP3^-/-^ mice at 3 and 7 days following TBI using FACs. The 3xTg mice start to show cognitive deficits around 6 months old, extracellular Aβ plaques around 9-12 months old and tau pathology around 12 months old. We injured the animals at the age around 4 months old before the onset of AD to enable us to examine injury-induced immune changes as a consequence of AD predisposition. Both male and female mice were included in the study and analyzed separately. Thus, the data comparison was made between the same sex 3xTg and 3xTg/NLRP3^-/-^ mice at the same post-injury time point.

The myeloid cells including infiltrating neutrophils (CD11b^+^Ly6G^+^), macrophages (CD45^high^CD11b^high^), and the brain residential microglia (CD45^low^CD11b^low^) were assessed. Among all animal groups, the baseline level of all three types of cells in the brain was comparable in the sham 3xTg and 3xTg/NLRP3^-/-^ mice regardless of sex. Following TBI, genotype and sex related differences were found among neutrophils and macrophages, but not microglia.

Specifically, at the acute stage following TBI (**3 dpi**), in female mice, a significant injury effect was observed in neutrophils and macrophages with a significantly higher number of these cells infiltrating the brain following TBI in both 3xTg and 3xTg/NLRP3^-/-^ mice when compared to their genotype matched sham controls (**Fig. 1A and B**, p<0.0001). When genotype comparison was made to assess the importance of NLRP3 in affecting the injury effect on these cells, only the number of infiltrating neutrophils was significant reduced in the injured 3xTg/NLRP3^-/-^ group in comparison to the injured 3xTg mice (**Fig. 1A**, p<0.0001). In male mice, TBI induced a higher number of brain-infiltrating neutrophils and macrophages in 3xTg mice similar to the 3xTg female mice (**Fig. 1F and G**, p<0.0001). The injury effect on these cells was diminished in male mice following NLRP3 gene deletion, as injured 3xTg/NLRP3^-/-^ mice had a significant reduction of both neutrophils and macrophages compared to the injured 3xTg mice when genotype comparison was made (**Fig. 1F and G**, p<0.0001). At 3 dpi, in both male and female mice, microglia did not show significant changes related to either injury or genotype (**Fig. 1C and H**).

**Fig. 1.**
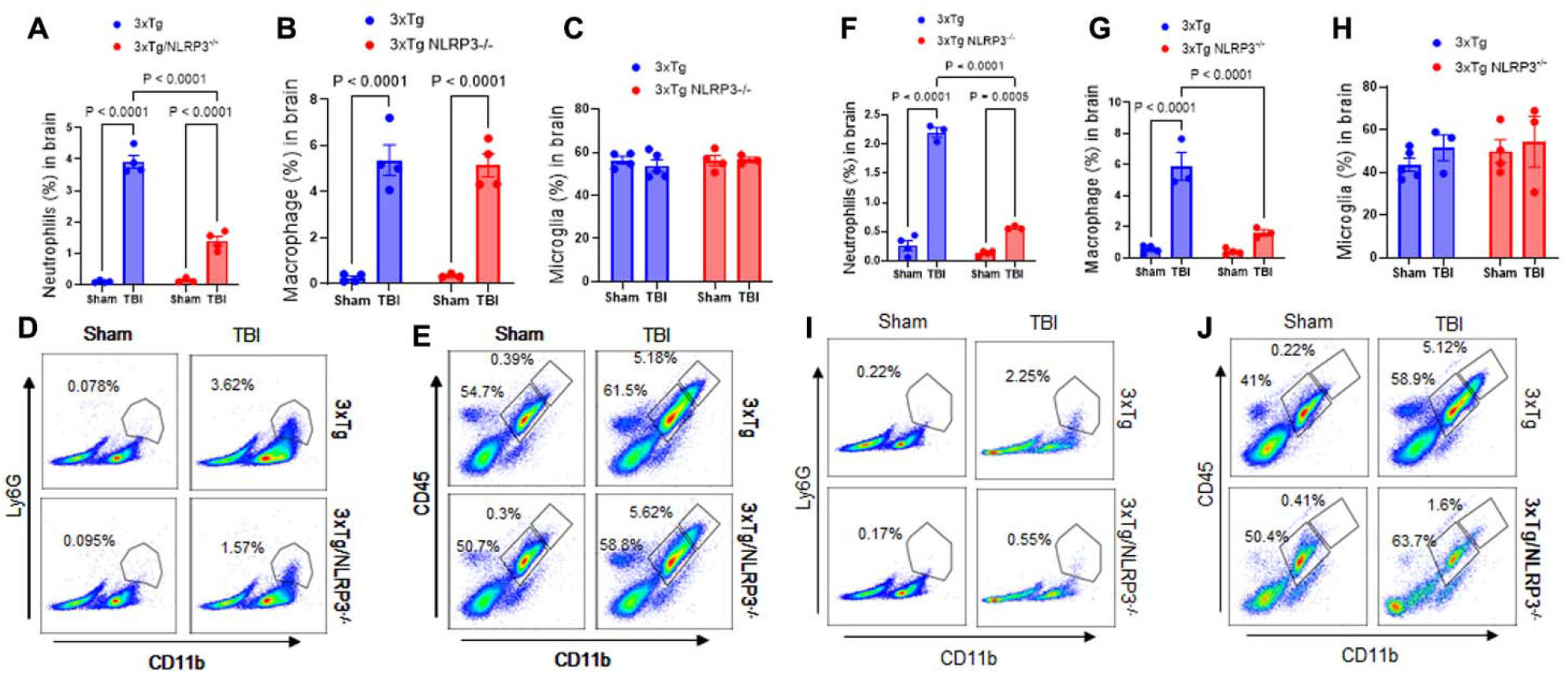
TBI-induced innate immune cell profiling changes in 3xTg or 3xTg/NLRP3-/-mice at acute stage (3dpi) following TBI. **(A-C and F-H):** Graphs show quantification analysis of frequencies of neutrophils (CD11b^+^Ly6G^+^), microglia (CD45^low^CD11b^low^) and infiltrating macrophages (CD45^high^CD11b^high^) infiltrating into the brain at 3dpi by FACs. **Female mice (A-E):** TBI significantly increased the number of neutrophils (**A**, p<0.0001) and macrophages (**B**, p<0.0001) in both 3xTg and 3xTg/NLRP3^-/-^ mice compared to their matched sham. Among the injured animals, 3xTg/NLRP3^-/-^ mice had significantly fewer infiltrating neutrophils compared to injured 3xTg group (A, p<0.0001). **Male mice (F-J):** TBI significantly increased the number of neutrophils (**F**, p<0.0001) and macrophages in 3xTg mice (**G**, p<0.0001) compared to matched sham, whereas in 3xTg/NLRP3^-/-^ mice, TBI only induced a moderate increase of neutrophils (**F**, p=0.0005) compared to matched sham. Among the injured animals, 3xTg/NLRP3^-/-^ mice had significantly fewer infiltrating neutrophils (**F**, p<0.0001) and macrophages (**G**, p<0.0001) compared to injured 3xTg mice. In both males and females, microglia did not show injury or genotype related changes (**C and H**). (**D-E and I-J)**. Representative gating images of infiltrating neutrophils (CD11b^+^Ly6G^+^), macrophages (CD45^high^CD11b^high^), and microglial (CD45^low^CD11b^low^) in the brain.

At the subacute stage following TBI (**7dpi**), in female mice, no difference was found between injured and sham groups in the number of all three types of cells in both 3xTg and 3xTg/NLRP3^-/-^ mice (**Fig. 2A-E**). In male mice, injury-induced macrophage infiltration remained high in the injured 3xTg mice only (**Fig. 2G**, p<0.0149), whereas no difference was found between injured and sham groups in the number of neutrophils and microglia in both 3xTg and 3xTg/NLRP3^-/-^ mice (**Fig. 2F and H**).

**Fig. 2.**
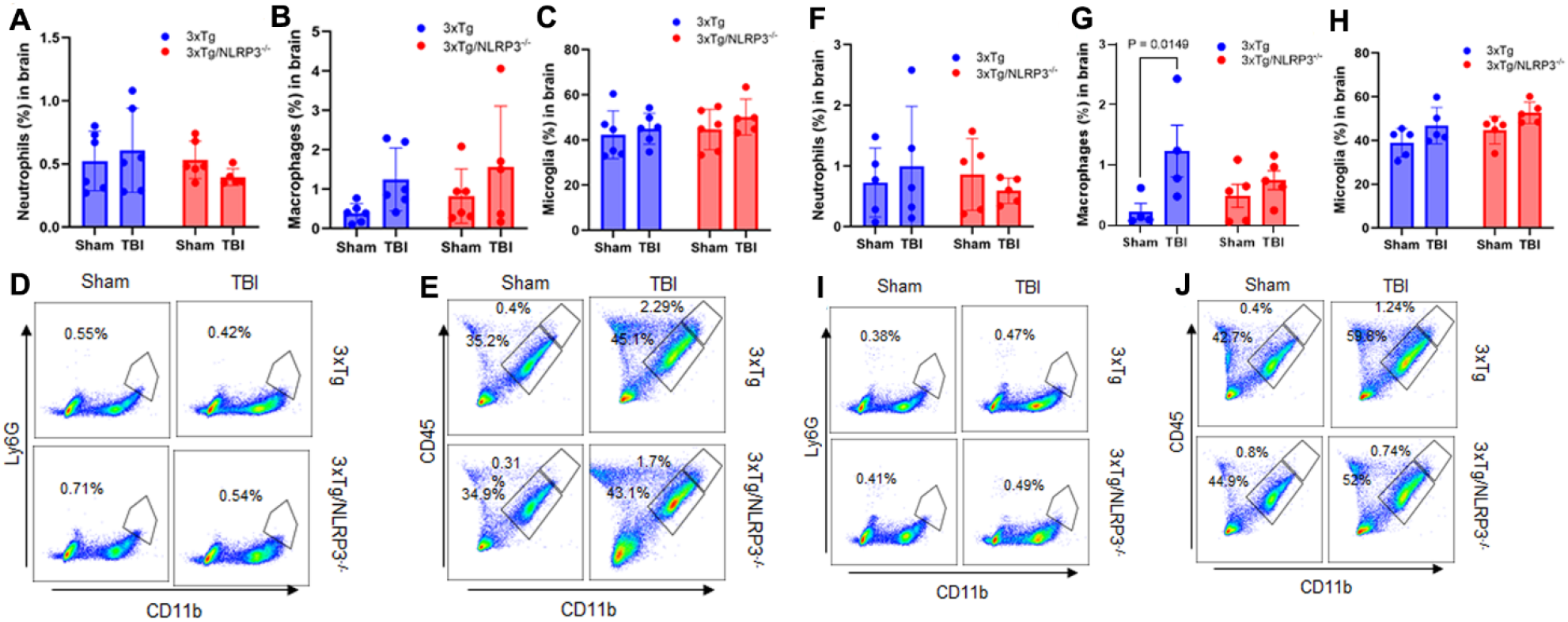
TBI-induced innate immune cell profiling changes in 3xTg or 3xTg/NLRP3^-/-^ mice at acute stage (7dpi) following TBI. **(A-C and F-H):** Graphs show quantification analysis of frequencies of neutrophils (CD11b^+^Ly6G^+^), microglia (CD45^low^CD11b^low^) and infiltrating macrophages (CD45^high^CD11b^high^) infiltrating into the brain at 7dpi by FACs. **Female mice (A-E)**: No significant injury induced change in the number of neutrophils (**A**), macrophages (**B**) and microglia (**C**) was observed in both 3xTg and 3xTg/NLRP3^-/-^ mice compared to their matched sham. **Male mice (F-J):** TBI-induced higher number of macrophages was observed in the injured brain in 3xTg mice (**G**, p=0.0149) compared to matched sham, but not in 3xTg/NLRP3^-/-^ mice. No injury or genotype related difference was found in the number of neutrophils (**F**) and microglia (**H**) in male mice. (**D-E and I-J)**. Representative gating images of infiltrating neutrophils (CD11b^+^Ly6G^+^), macrophages (CD45^high^CD11b^high^), and microglial (CD45^low^CD11b^low^) in the brain.

### TBI induces changes in adaptive immunity in 3xTg mice at the acute stage following injury and loss of NLRP3 counteracts the injury effect

We also assessed the effect of TBI and NLRP3 gene deletion on responses of IL-17^+^ helper T cells (CD4^+^IL-17A) and gamma delta T cells (γδ T cells, γδTCR^+^IL-17A^+^). Gamma delta T cells (γδ T cells) are considered as regulatory T cells bridging innate and adaptive immune responses, and their accumulation in the brain indicates that they act as crucial regulators of brain inflammation. In our study, an injury and NLRP3 null induced γδ T cell response was observed at the acute stage following TBI. Specifically, at 3 dpi, higher numbers of γδ T cells were observed in the brains of injured 3xTg mice in both males (**Fig. 3A**, p<0.0001), and females (**Fig. 3B**, p<0.0001) when compared to the matched sham, while this injury effect was not observed following NLRP3 gene deletion in both sexes (**Fig. 3A and B**). When genotype comparison was made, injured 3xTg/NLRP3^-/-^ mice had significantly lower frequency of γδ T cells in the brain compared to injured 3xTg mice in both sexes (**Fig. 3A and B**, male: p=0.0002; female: p=0031). For CD4 cells, at 3 dpi, higher numbers of infiltrating IL-17^+^ CD4 cells were found in the injured brain only in female 3xTg and 3xTg/NLRP3^-/-^ mice compared to their genotype matched sham (**Fig. 3D**), not in male mice (**Fig. 3C**). At 7dpi, no difference was observed in the frequency of γδ T and IL-17^+^ CD4 cells among all groups studied (**Supplemental Fig. 2**).

**Fig. 3.**
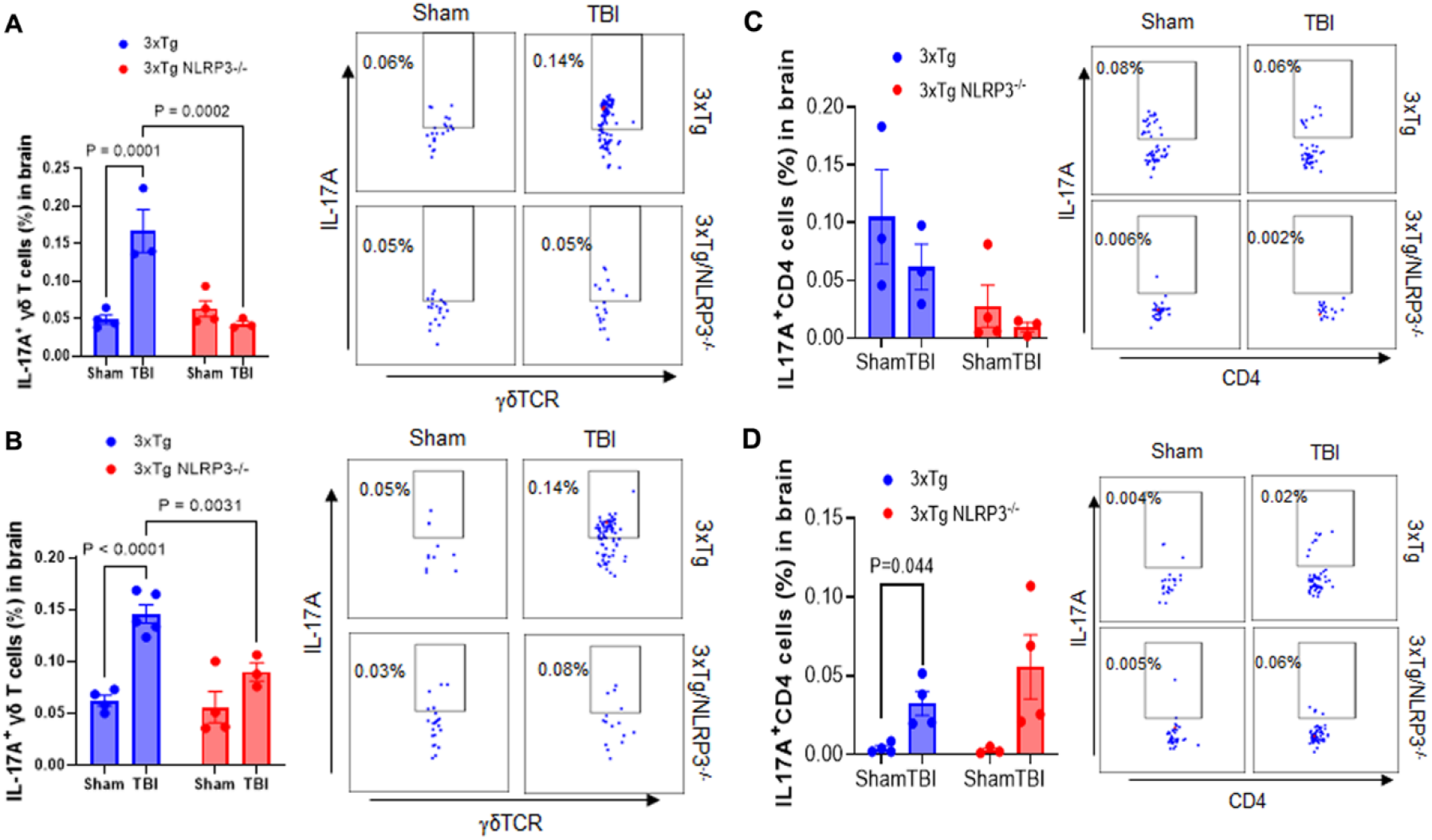
TBI-induced infiltration of γδT and IL-17^+^ CD4 cells in the brain at 3 dpi. Frequencies of activated γδT cells (γδTCR^+^IL-17A^+^) and IL-17^+^ CD4 cells (CD4^+^IL-17A^+^) infiltrating into the brain were analyzed by flow cytometry. **A-B**). Significant higher number of γδT cells was observed in the injured 3xTg mice in both males (**A**) and females (**B**) as compared to their matched sham, while this injury effect was not observed in the 3xTg/NLRP3^-/-^ mice in both sexes. Moreover, significantly lower number of γδT cells was found in the injured 3xTg/NLRP3^-/-^ mice in comparison to the injured 3xTg mice in both sexes. **C-D**). TBI increased significant infiltration of IL-17^+^ CD4 cells in female 3xTg mice (**D**) but not in male 3xTg mice (**C**), and NLRP3 deletion did not show significant impact.

### Deletion of NLRP3 gene reduces TBI-enhanced NLRP3 downstream cytokine gene expression only in male AD mice with opposite effect in female mice

Using real-time PCR, we assessed the influence of TBI and the NLRP3 inflammasome on gene expression levels of NLRP3 downstream targets including IL-1β, TNF-α, IL-18 and IL-17f. A sex-related differential expression pattern was found in response to injury and NLRP3 deletion.

At the acute stage following TBI (**3 dpi**), in female mice, the expression level of IL-1β, TNF-α, IL-18 and IL-17f genes was not affected by injury in the 3xTg mice (**Fig. 4A-D**). In contrast, a significant injury effect was found in 3xTg/NLRP3^-/-^ mice which had increased expression of IL-1β and TNF-α genes in the injured animals when compared to the matched sham (**Fig. 4A/B**, IL-1β: p=0.007; TNF-α: p<0.0001), and the injured 3xTg mice (**Fig. 4A/B**, IL-1β: p=0.0049; TNF-α: p<0.0001). A lower level of IL-18 expression was found in the female 3xTg/NLRP3^-/-^ mice than the 3xTg mice (**Fig. 4C**) though. In male mice, an opposite response was found with TBI inducing a significant increase of IL-1β and TNF-α gene expression in the 3xTg mice **(Fig. 4E/F**, IL-1β: p=0.003; TNF-α: p=0.005), but not in the 3xTg/NLRP3^-/-^ mice when compared to their genotype matched sham. Comparing to the injured 3xTg mice, the injured 3xTg/ NLRP3^-/-^ mice had a significantly lower level of IL-1β and TNF-α genes (**Fig. 4E/F**, IL-1β: p=0.047; TNF-α: p=0.0254). The expression level of IL-18 and IL-17f showed no change in the 3xTg mice following injury, however, was lower in the 3xTg/NLRP3^-/-^ mice (**Fig. 4G/H**)

**Fig. 4.**
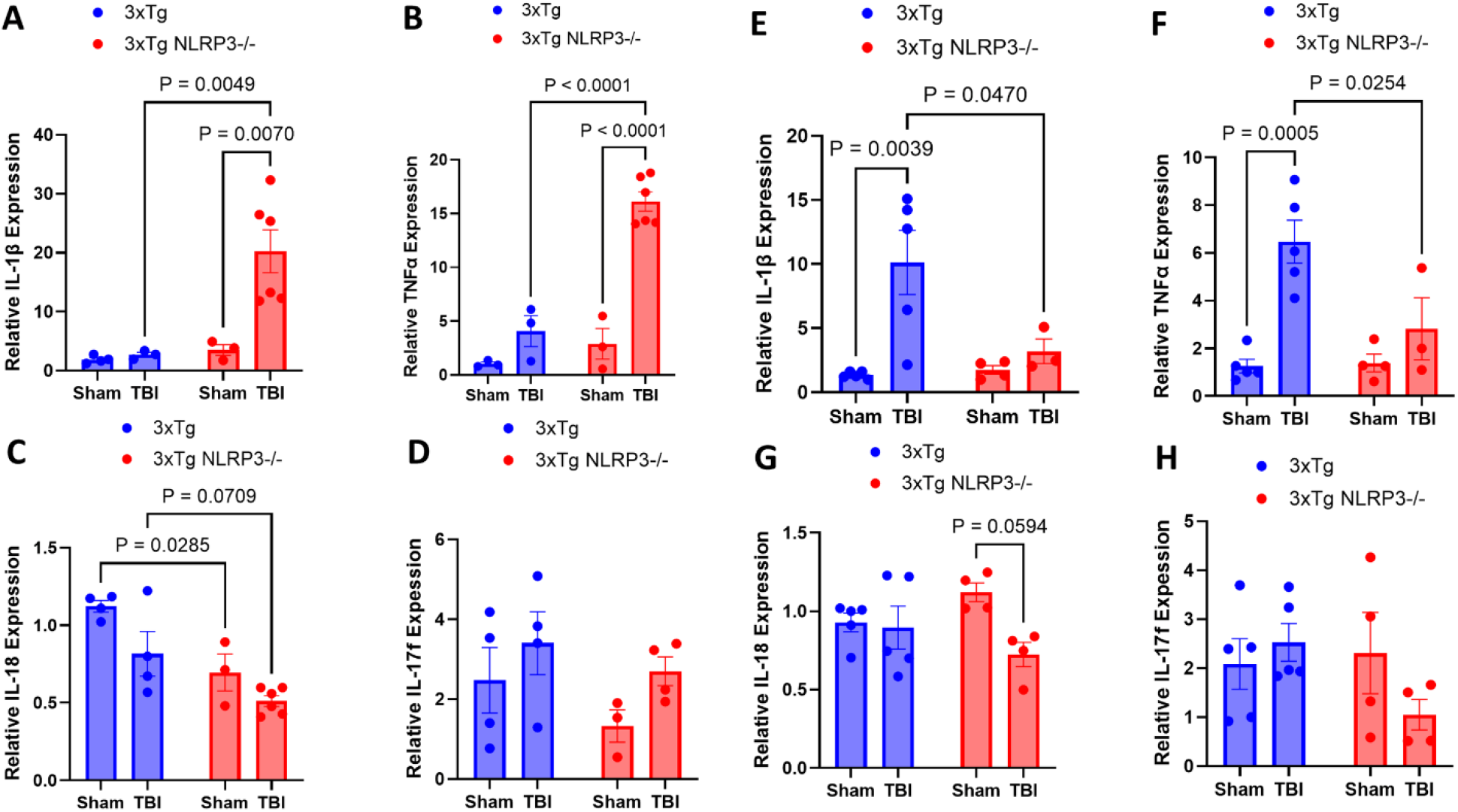
TBI-induced inflammatory gene expression changes in male and female mice at 3 dpi. Gene transcription of IL-1β, TNF-α, IL-18 and IL-17f in the brain was determined by real-time PCR. **A-D**). **Female mice:** In the 3xTg mice, TBI did not increase gene expression of IL-1β, TNF-α, IL-18 and IL-17f. In the 3xTg/NLRP3^-/-^ mice, significantly higher expression of IL-1β (**A**) and TNF-α (**B**) was found compared to matched sham and the injured 3xTg mice, while lower expression of IL-18 (**C**) was found in the 3xTg/NLRP3^-/-^ mice compared to the 3xTg mice and no change in IL-17f expression (**D**). **E-H**). **Male mice:** In the 3xTg mice, TBI significantly increased expression of IL-1β (**E**) and TNF-α (**F**) in the injured 3xTg mice compared to the matched sham. This injury effect was not observed in the 3xTg/NLRP3^-/-^ mice, with significantly lower expression of IL-1β, TNF-α in the injured 3xTg/NLRP3^-/-^ mice compared to the injured 3xTg mice. The expression level of IL-18 was not affected by injury in the 3xTg mice but lower expression was found in the injured 3xTg/NLRP3^-/-^ mice compared to matched sham (**G**). The expression level of IL-17f (H) was not affected by injury in both 3xTg and 3xTg/NLRP3^-/-^ mice (**H**).

At the subacute stage following TBI (**7 dpi**), injury induced elevation of IL-1β and TNFα genes remained high in the injured male 3xTg mice when compared to the matched sham (**Fig. 5A**). In the injured 3xTg/NLRP3^-/-^ mice, IL-1β gene expression was higher than the matched sham (**Fig. 5A**, p=0.0063), but was significantly lower than the injured 3xTg mice (**Fig. 5A**, p=0.0128), while TNFα was not significantly affected (**Fig. 5A**). In female mice, injury-enhanced elevation of IL-1β and TNFα genes remained similarly high in both 3xTg and 3xTg/NLRP3^-/-^ mice when compared to the matched sham (**Fig. 5B**, IL-1β: 3xTg, p=0.0004; 3xTg/NLRP3^-/-^, p=0.0002. TNFα: 3xTg, p=0.0027; 3xTg/NLRP3^-/-^, p=0.0011). Similar to 3dpi, no injury effect was found in the expression levels of IL-18 and IL-17f in 3xTg mice in both sexes. However, a slight reduction in IL-18 levels was found in the male injured 3xTg/NLRP3^-/-^ mice when compared to matched sham (**Fig. 5A**, p=0.0325), and the female injured 3xTg/NLRP3^-/-^ mice when compared to female injured 3xTg mice (**Fig. 5B**, p=0.0333).

**Fig. 5.**
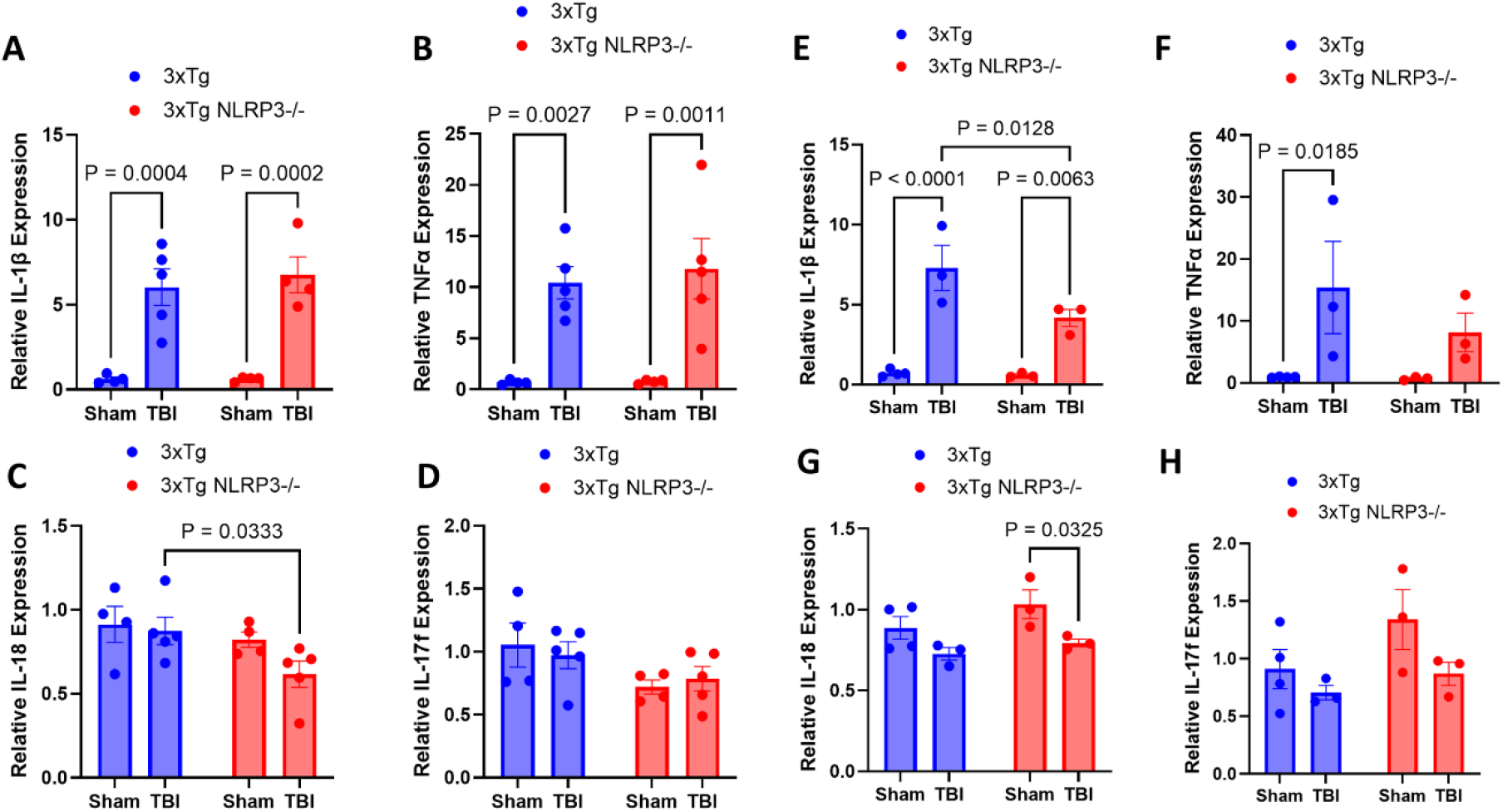
TBI-induced inflammatory gene expression changes in male and female mice at 7 dpi. Gene transcription of IL-1β, TNF-α, IL-18 and IL-17f in the brain was determined by real-time PCR. **A-D**). **Female mice:** TBI induced a comparable increase in IL-1β (**A**) and TNF-α (**B**) gene expression in both the 3xTg and 3xTg/NLRP3^-/-^ mice. No injury effect was found in expression level of IL-18 (**C**) and IL-17f (**D**) in both the 3xTg and 3xTg/NLRP3^-/-^ mice, however, lower expression of IL-18 (**C**) was found in the injured 3xTg/NLRP3^-/-^ mice compared to the injured 3xTg mice. **E-H**). **Male mice:** In the 3xTg mice, TBI significantly increased the expression level of IL-1β (**E**) and TNF-α (**F**). In the 3xTg/NLRP3^-/-^ mice, only IL-1β was elevated following TBI and the level was significantly lower than the injured 3xTg group (**E**). The injured 3xTg/NLRP3^-/-^ mice also had lower IL-18 expression than the matched sham (**G**), while the expression level of IL-17f remained unchanged in both the 3xTg and 3xTg/NLRP3^-/-^ mice (**H**).

The PCR data suggested a sustained injury-effect on neuroinflammation in the 3xTg mice unrelated to the immune cell profiling. Furthermore, the female mice exhibited an NLRP3 independent inflammatory response following TBI.

## Discussion

TBI is a major risk factor for AD, however, the mechanistic link between the two is unclear. Extensive studies have implicated that NLRP3 inflammasome plays an important role in AD, while TBI activates NLRP3 inflammasome. In this study, we investigated the importance of the NLRP3 inflammasome in regulating the post-TBI immune/inflammatory response in the context of AD predisposition using a novel 3xTg/NLRP3 null mouse line. We also compared sex-related differences in relation to TBI and AD. We found that TBI altered immune profiles in 3xTg mice while NLRP3 gene knock out counteracts the injury effect with a significant sex-related difference. More specifically, at the acute stage after TBI in both male and female mice, TBI induced a significant infiltration of neutrophils, macrophages and γδ T-cells in the brains of 3xTg mice, whereas NLRP3 knock down significantly blocked this injury effect in males but less so in females. We also found that NLRP3 gene deletion counteracted TBI-enhanced expression of inflammatory cytokines IL-1β, TNF-α, and IL-17F. In summary, this study has confirmed that TBI significantly alters immune profiles in the context of AD predisposition, the NLRP3 inflammasome is important in mediating these TBI-induced changes, and there are significant sex-related differences.

Neuroinflammation is an essential player driving disease development in many neurological disorders including TBI and AD. In AD pathogenesis, inflammation is an important player, and TBI significantly exacerbates this response. Studies have reported that even a single mild injury induces an abnormal inflammatory response exacerbating AD pathology and cognitive deficits in APP/PS1 mice (Webster et al., 2015). The effects of the immune response as a result of injury are well documented with many contributing proteins and signaling molecules identified and evaluated. NLRP3 is a more recent protein with an integral role in the inflammatory and immune response to injury or infection. In Alzheimer’s disease, NLRP3 is known to play an integral role in the propagation of the disease and promote tau pathology (Heneka et al., 2013). Additionally, in the brain, NLRP3 is one of the sources of inflammation that originates directly from the microglia (Freeman et al., 2017; Liu et al., 2020). Following TBI, upregulation of the NLRP3 inflammasome was observed in microglia, astrocytes and vascular endothelial cells contributing to neuroinflammation via the release of IL-1β and pyroptosis in neurons (Irrera et al., 2020; O’Brien et al., 2020; Shaheen et al., 2021; Thakkar et al., 2016; Yi et al., 2020; Zhang et al., 2025). In the present study, we assessed the direct effects of NLRP3 in mediating TBI-induced innate and adaptive immune responses in the context of predisposition of AD via a genetic knockout of NLRP3 gene in 3xTg mice. By immune cell profiles with FACs and PCR measurements of cytokine gene expression at the acute and subacute stages following TBI, we have found that in the context of predisposition of AD, there are significant immune changes including both innate (neutrophils and macrophages) and adaptive immune (CD4 and γδ T cells) responses in the brain following a moderate TBI. Neutrophils are an abundant immune cell found in the blood that are able to penetrate into the brain upon injury. In TBI, neutrophils peak in abundance around day 2 and decrease in number until about day 7 post injury (Liu et al., 2018). In the context of TBI, neutrophils are believed to influence blood flow, cerebrospinal fluid levels, BBB integrity, hypoxia, neuroinflammation, neurodegeneration, and neuronal recovery (Liu et al., 2018). Our results confirm the timing of injury-induced neutrophil infiltration into the brain. Our study also suggests a regulatory role of NLRP3 in this process as both male and female 3xTg/NLRP3^-/-^ mice have reduced levels of neutrophil infiltration following TBI compared to 3xTg mice (**Fig. 1A** and **F**). Macrophages and microglia play a similar critical role in the immune response to TBI. However, they differ in the response to damage-associated molecular patterns (DAMPs) and other innate immune responses to promote immune cell recruitment and inflammation via NLRP3 (Zarruk et al., 2018). Our results have found injury-enhanced infiltration of macrophages into the brain at the acute stage following TBI in 3xTg mice in both male and females (**Fig. 1B and G**), and it persists to the subacute stage in males but not in females (**Fig. 2B and G**). NLRP3 gene deletion blocked the injury effect only in males not in females (**Fig. 1B and G)**. Our data suggests that the TBI-induced infiltration of macrophages is transient at a more heightened level in females, whereas in males this response lasts longer. At both the acute and subacute stages in both sexes regardless of whether NLRP3 was present or absent, microglia cell numbers exhibited no significant injury-induced changes in our animals (**Fig. 2-3C and H**), which is in agreement with other studies reporting that infiltrating myeloid cells were significantly more reactive than resident microglia following TBI (Doran et al., 2019).

In the current study, we have also examined the response of γδ T cells to injury and NLRP3 knock down. As the bridge of innate and adaptive immunities, γδ T cells enriched in the brain are crucial regulators of CNS inflammation (Dressman and Elyaman, 2022; Li et al., 2021; Wo et al., 2020). Production of IL-17A by γδT cells in the absence of αβT cells is critical for the initiation of CNS inflammation. However, the role of γδT cells in TBI and AD remains to be elucidated. It is known that IL-1β, IL-18, and IL-23 can promote the secretion of IL-17A from γδT cells in the absence of primary (TCR) and secondary (co-stimulator) signals(Lalor et al., 2011; Sutton et al., 2009). It has also been demonstrated that IL-17A can further recruit IL-1β-secreting myeloid cells amplifying neuroinflammation (Lalor et al., 2011). Thus, the cooperative actions of resident and infiltrating immune cells (microglial/macrophages/monocytes, γδ T cells) promote pathogenic neuroinflammation in CNS diseases, including TBI and AD(Dressman and Elyaman, 2022; Kokiko-Cochran and Godbout, 2018; Wo et al., 2020). In the current study, we have found significant injury-induced activation of γδ T cells in 3xTg mice but not 3xTg/NLRP3^-/-^ mice in both males and females in the acute stage following TBI (**Fig. 3A and B**). This observation for the first time establishes the potential association between NLRP3 inflammasome activity and activation of IL-17-producing γδ T cells following TBI.

In the current study, we have examined the influence of injury and NLRP3 on the immune response in both males and females, as there is a significant sex-related difference in both TBI and AD (Doran et al., 2019; Royal et al., 2026; Villapol et al., 2017). Thus far published studies examining NLRP3 in the context of TBI have primarily been performed in males (O’Brien et al., 2020). Here we report notable changes in innate immune responses to TBI in males and females. Apart from injury-induced differences in the degree of infiltration of neutrophils and macrophages, the expression level of pro-inflammatory cytokine IL-1β and TNF-α also showed significant sex-related differences. In males, injury-enhanced expression of IL-1β and TNF-α were found in both 3 and 7 dpi in 3xTg males, while females showed injury-induced upregulation at 7 dpi but not at 3 dpi (**Fig. 4**). This sex-related difference in post-TBI cytokine gene expression pattern in 3xTg mice is similar to others reported in wild type mice (Villapol et al., 2017). Uniquely in our 3xTg/NLRP3^-/-^ mice, deletion of NLRP3 gene blocked injury-enhanced expression of IL-1β and TNF-α in male but not female animals suggesting a sex-related difference in regulating cytokine expression following TBI.

While our results clearly show a connection between NLRP3 and the immune cell response following TBI, we did not analyze the inflammasome components directly nor what specifically might be causing these changes. Further studies would need to look into expanding the reasons for the sex related differences we observed as well as how NLRP3 is specifically affected by sex hormonal signaling. Additionally, we have only reported results analyzing the acute and subacute stages following injury. Studying the long-term influence of NLRP3 on the post-TBI chronic immune/inflammatory response and if that leads to AD pathological development warrant further examination. Investigating the effects of TBI on advanced aged mice likewise warrant further investigation as older individuals clinically account for a significant portion of all TBI patients.

## Acknowledgements

This work was supported by Commonwealth Health Research Board grant (#236-12-25, DS), Commonwealth of Virginia Center on Aging - Alzheimer’s and Related Disease Research Award Fund (#21-1, DS), NIA/NIH (U01AG076481, Zhang). Services and products in support of the research project were generated by the Virginia Commonwealth University Flow Cytometry Shared Resource, supported, in part, with funding from NIH-NCI Cancer Center Support Grant P30 CA016059.

## Declarations

### Availability of data and materials

Data generated or analyzed for this study are available from the corresponding author on reasonable request.

### Authors’ contribution

JG was responsible for FACs study, statistical analysis, and wrote the draft of manuscript. ZL and CG performed FACs and PCR experiments and statistical analysis. ST and CN performed TBI. AP performed WB analysis. XW assisted FACs and PCR data interpretation and manuscript editing. SZ assisted with funding acquisition and manuscript editing. DS conceived the conceptual idea, designed the study, and was responsible for funding acquisition and manuscript writing.

### Generative AI use

No AI and AI-assisted tools were used during the preparation of this work.

### Ethical approval and Consent to participate

No human tissue was involved in this study. Animal study was performed strictly following guidelines of IACUC, all procedures were approved by IACUC.

### Consent for publication

The submission was approved by all authors.

## Sumpplemental Data

**Suppl-Fig. 1.**
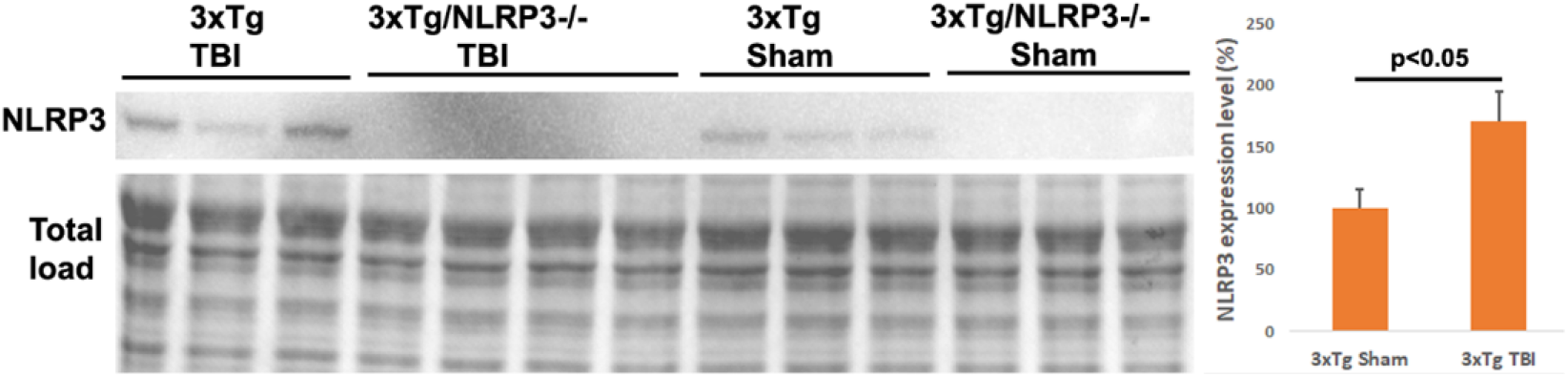
Protein expression level of NLRP3 is increased following TBI in 3xTg mice and is absent in 3xTg/NLRP3^-/-^ mice. Western blotting image shows expression of NLRP3 protein (120KDa) in the cerebral cortex in male mice at 3 days post-TBI. Increased NLRP3 expression was observed in the injured 3xTg mice compared to sham while it was absent in 3xTg/NLRP3^-/-^ mice.

**Suppl-Fig. 2.**
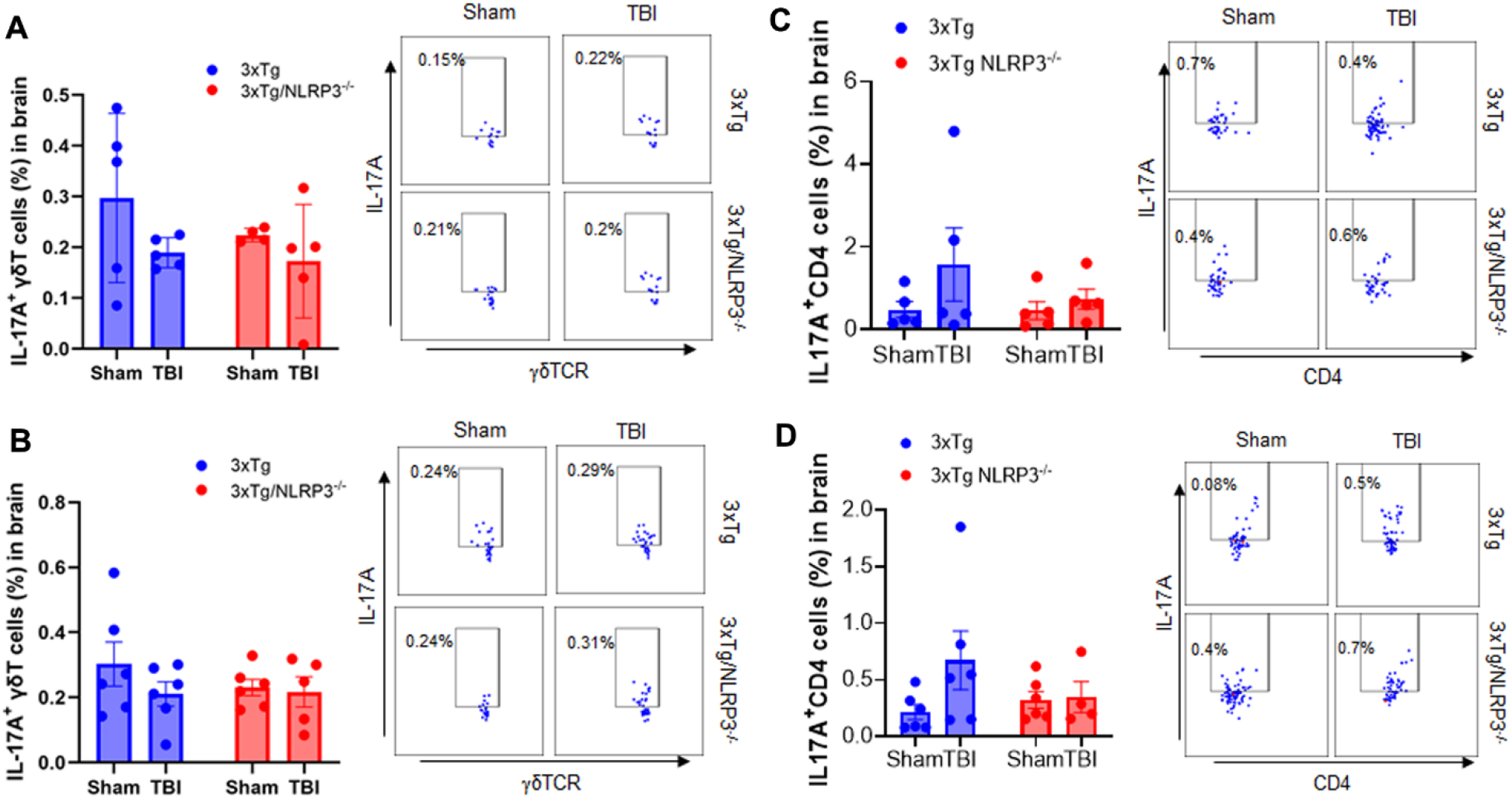
TBI-induced infiltration of γδT and IL-17^+^ CD4 cells in the brain at 7 dpi. Frequencies of activated γδT cells (γδTCR^+^IL-17A^+^, **A-B**) and IL-17^+^ CD4 cells (CD4^+^IL-17A^+^, **C**-**D)** infiltrating into the brain were analyzed by flow cytometry. No significant injury or genotype-induced difference in the number of infiltrating γδT cells (**A-B**), or IL-17^+^ CD4 cells (**C-D**) was found in both male (**A, C**) and female mice (**B, D**).

## References

Adamczak, S., G. Dale, J.P. de Rivero Vaccari, M.R. Bullock, W.D. Dietrich, and R.W. Keane. 2012. Inflammasome proteins in cerebrospinal fluid of brain-injured patients as biomarkers of functional outcome: clinical article. J Neurosurg. 117:1119–1125.

Adamczak, S.E., J.P. de Rivero Vaccari, G. Dale, F.J. Brand, 3rd, D. Nonner, M.R. Bullock, G.P. Dahl, W.D. Dietrich, and R.W. Keane. 2014. Pyroptotic neuronal cell death mediated by the AIM2 inflammasome. J Cereb Blood Flow Metab. 34:621–629.

Byers, S.L., M.V. Wiles, S.L. Dunn, and R.A. Taft. 2012. Mouse estrous cycle identification tool and images. PLoS One. 7:e35538.

Couturier, J., I.C. Stancu, O. Schakman, N. Pierrot, F. Huaux, P. Kienlen-Campard, I. Dewachter, and J.N. Octave. 2016. Activation of phagocytic activity in astrocytes by reduced expression of the inflammasome component ASC and its implication in a mouse model of Alzheimer disease. J Neuroinflammation. 13:20.

Cumiskey, D., M. Pickering, and J.J. O’Connor. 2007. Interleukin-18 mediated inhibition of LTP in the rat dentate gyrus is attenuated in the presence of mGluR antagonists. Neurosci Lett. 412:206–210.

Doran, S.J., R.M. Ritzel, E.P. Glaser, R.J. Henry, A.I. Faden, and D.J. Loane. 2019. Sex Differences in Acute Neuroinflammation after Experimental Traumatic Brain Injury Are Mediated by Infiltrating Myeloid Cells. J Neurotrauma. 36:1040–1053.

Dressman, D., and W. Elyaman. 2022. T Cells: A Growing Universe of Roles in Neurodegenerative Diseases. Neuroscientist. 28:335–348.

Faul, M., and V. Coronado. 2015. Epidemiology of traumatic brain injury. Handb Clin Neurol. 127:3–13.

Freeman, L., H. Guo, C.N. David, W.J. Brickey, S. Jha, and J.P. Ting. 2017. NLR members NLRC4 and NLRP3 mediate sterile inflammasome activation in microglia and astrocytes. J Exp Med. 214:1351–1370.

Freeman, L.C., and J.P. Ting. 2016. The pathogenic role of the inflammasome in neurodegenerative diseases. J Neurochem. 136 Suppl 1:29–38.

Gardner, R.C., J.F. Burke, J. Nettiksimmons, A. Kaup, D.E. Barnes, and K. Yaffe. 2014. Dementia risk after traumatic brain injury vs nonbrain trauma: the role of age and severity. JAMA Neurol. 71:1490–1497.

Gardner, R.C., and K. Yaffe. 2014. Traumatic brain injury may increase risk of young onset dementia. Ann Neurol. 75:339–341.

Ghosh, S., M.D. Wu, S.S. Shaftel, S. Kyrkanides, F.M. LaFerla, J.A. Olschowka, and M.K. O’Banion. 2013. Sustained interleukin-1beta overexpression exacerbates tau pathology despite reduced amyloid burden in an Alzheimer’s mouse model. J Neurosci. 33:5053–5064.

Griffin, W.S., L.C. Stanley, C. Ling, L. White, V. MacLeod, L.J. Perrot, C.L. White, 3rd, and C. Araoz. 1989. Brain interleukin 1 and S-100 immunoreactivity are elevated in Down syndrome and Alzheimer disease. Proc Natl Acad Sci U S A. 86:7611–7615.

Gris, D., Z. Ye, H.A. Iocca, H. Wen, R.R. Craven, P. Gris, M. Huang, M. Schneider, S.D. Miller, and J.P. Ting. 2010. NLRP3 plays a critical role in the development of experimental autoimmune encephalomyelitis by mediating Th1 and Th17 responses. J Immunol. 185:974–981.

Halle, A., V. Hornung, G.C. Petzold, C.R. Stewart, B.G. Monks, T. Reinheckel, K.A. Fitzgerald, E. Latz, K.J. Moore, and D.T. Golenbock. 2008. The NALP3 inflammasome is involved in the innate immune response to amyloid-beta. Nat Immunol. 9:857–865.

Hellewell, S., B.D. Semple, and M.C. Morganti-Kossmann. 2016. Therapies negating neuroinflammation after brain trauma. Brain Res. 1640:36–56.

Heneka, M.T. 2017. Inflammasome activation and innate immunity in Alzheimer’s disease. Brain Pathol. 27:220–222.

Heneka, M.T., M.J. Carson, J. El Khoury, G.E. Landreth, F. Brosseron, D.L. Feinstein, A.H. Jacobs, T. Wyss-Coray, J. Vitorica, R.M. Ransohoff, K. Herrup, S.A. Frautschy, B. Finsen, G.C. Brown, A. Verkhratsky, K. Yamanaka, J. Koistinaho, E. Latz, A. Halle, G.C. Petzold, T. Town, D. Morgan, M.L. Shinohara, V.H. Perry, C. Holmes, N.G. Bazan, D.J. Brooks, S. Hunot, B. Joseph, N. Deigendesch, O. Garaschuk, E. Boddeke, C.A. Dinarello, J.C. Breitner, G.M. Cole, D.T. Golenbock, and M.P. Kummer. 2015. Neuroinflammation in Alzheimer’s disease. Lancet Neurol. 14:388–405.

Heneka, M.T., M.P. Kummer, A. Stutz, A. Delekate, S. Schwartz, A. Vieira-Saecker, A. Griep, D. Axt, A. Remus, T.C. Tzeng, E. Gelpi, A. Halle, M. Korte, E. Latz, and D.T. Golenbock. 2013. NLRP3 is activated in Alzheimer’s disease and contributes to pathology in APP/PS1 mice. Nature. 493:674–678.

Iacono, D., S. Raiciulescu, C. Olsen, and D.P. Perl. 2021. Traumatic Brain Injury Exposure Lowers Age of Cognitive Decline in AD and Non-AD Conditions. Front Neurol. 12:573401.

Irrera, N., G. Pizzino, M. Calo, G. Pallio, F. Mannino, F. Fama, V. Arcoraci, V. Fodale, A. David, C. Francesca, L. Minutoli, E. Mazzon, P. Bramanti, F. Squadrito, D. Altavilla, and A. Bitto. 2017. Lack of the Nlrp3 Inflammasome Improves Mice Recovery Following Traumatic Brain Injury. Front Pharmacol. 8:459.

Irrera, N., M. Russo, G. Pallio, A. Bitto, F. Mannino, L. Minutoli, D. Altavilla, and F. Squadrito. 2020. The Role of NLRP3 Inflammasome in the Pathogenesis of Traumatic Brain Injury. Int J Mol Sci. 21.

Ising, C., C. Venegas, S. Zhang, H. Scheiblich, S.V. Schmidt, A. Vieira-Saecker, S. Schwartz, S. Albasset, R.M. McManus, D. Tejera, A. Griep, F. Santarelli, F. Brosseron, S. Opitz, J. Stunden, M. Merten, R. Kayed, D.T. Golenbock, D. Blum, E. Latz, L. Buee, and M.T. Heneka. 2019. NLRP3 inflammasome activation drives tau pathology. Nature. 575:669–673.

Johnson, V.E., W. Stewart, and D.H. Smith. 2012. Widespread tau and amyloid-beta pathology many years after a single traumatic brain injury in humans. Brain Pathol. 22:142–149.

Julien, J., S. Joubert, M.C. Ferland, L.C. Frenette, M.M. Boudreau-Duhaime, L. Malo-Veronneau, and E. de Guise. 2017. Association of traumatic brain injury and Alzheimer disease onset: A systematic review. Ann Phys Rehabil Med. 60:347–356.

Kokiko-Cochran, O.N., and J.P. Godbout. 2018. The Inflammatory Continuum of Traumatic Brain Injury and Alzheimer’s Disease. Front Immunol. 9:672.

Krstic, D., and I. Knuesel. 2013. Deciphering the mechanism underlying late-onset Alzheimer disease. Nat Rev Neurol. 9:25–34.

Kumar, A., and D.J. Loane. 2012. Neuroinflammation after traumatic brain injury: opportunities for therapeutic intervention. Brain Behav Immun. 26:1191–1201.

Kuwar, R., A. Rolfe, L. Di, H. Xu, L. He, Y. Jiang, S. Zhang, and D. Sun. 2019. A novel small molecular NLRP3 inflammasome inhibitor alleviates neuroinflammatory response following traumatic brain injury. J Neuroinflammation. 16:81.

Lalor, S.J., L.S. Dungan, C.E. Sutton, S.A. Basdeo, J.M. Fletcher, and K.H. Mills. 2011. Caspase-1-processed cytokines IL-1beta and IL-18 promote IL-17 production by gammadelta and CD4 T cells that mediate autoimmunity. J Immunol. 186:5738–5748.

Latz, E., T.S. Xiao, and A. Stutz. 2013. Activation and regulation of the inflammasomes. Nat Rev Immunol. 13:397–411.

Lee, D.C., J. Rizer, M.L. Selenica, P. Reid, C. Kraft, A. Johnson, L. Blair, M.N. Gordon, C.A. Dickey, and D. Morgan. 2010. LPS-induced inflammation exacerbates phospho-tau pathology in rTg4510 mice. J Neuroinflammation. 7:56.

Li, Y., Y. Zhang, and X. Zeng. 2021. gammadelta T Cells Participating in Nervous Systems: A Story of Jekyll and Hyde. Front Immunol. 12:656097.

Lin, C., H. Chao, Z. Li, X. Xu, Y. Liu, Z. Bao, L. Hou, Y. Liu, X. Wang, Y. You, N. Liu, and J. Ji. 2017. Omega-3 fatty acids regulate NLRP3 inflammasome activation and prevent behavior deficits after traumatic brain injury. Exp Neurol. 290:115–122.

Liu, H.D., W. Li, Z.R. Chen, Y.C. Hu, D.D. Zhang, W. Shen, M.L. Zhou, L. Zhu, and C.H. Hang. 2013. Expression of the NLRP3 inflammasome in cerebral cortex after traumatic brain injury in a rat model. Neurochem Res. 38:2072–2083.

Liu, Y., Y. Dai, Q. Li, C. Chen, H. Chen, Y. Song, F. Hua, and Z. Zhang. 2020. Beta-amyloid activates NLRP3 inflammasome via TLR4 in mouse microglia. Neurosci Lett. 736:135279.

Liu, Y.W., S. Li, and S.S. Dai. 2018. Neutrophils in traumatic brain injury (TBI): friend or foe? J Neuroinflammation. 15:146.

Loane, D.J., and K.R. Byrnes. 2010. Role of microglia in neurotrauma. Neurotherapeutics. 7:366–377.

Loane, D.J., and A.I. Faden. 2010. Neuroprotection for traumatic brain injury: translational challenges and emerging therapeutic strategies. Trends Pharmacol Sci. 31:596–604.

LoBue, C., H. Wadsworth, K. Wilmoth, M. Clem, J. Hart, Jr., K.B. Womack, N. Didehbani, L.H. Lacritz, H.C. Rossetti, and C.M. Cullum. 2017. Traumatic brain injury history is associated with earlier age of onset of Alzheimer disease. Clin Neuropsychol. 31:85–98.

Lozano, D., G.S. Gonzales-Portillo, S. Acosta, I. de la Pena, N. Tajiri, Y. Kaneko, and C.V. Borlongan. 2015. Neuroinflammatory responses to traumatic brain injury: etiology, clinical consequences, and therapeutic opportunities. Neuropsychiatr Dis Treat. 11:97–106.

Lynch, M.A. 2015. Neuroinflammatory changes negatively impact on LTP: A focus on IL-1beta. Brain Res. 1621:197–204.

Ma, M.W., J. Wang, K.M. Dhandapani, and D.W. Brann. 2017. NADPH Oxidase 2 Regulates NLRP3 Inflammasome Activation in the Brain after Traumatic Brain Injury. Oxid Med Cell Longev. 2017:6057609.

Mariathasan, S., D.S. Weiss, K. Newton, J. McBride, K. O’Rourke, M. Roose-Girma, W.P. Lee, Y. Weinrauch, D.M. Monack, and V.M. Dixit. 2006. Cryopyrin activates the inflammasome in response to toxins and ATP. Nature. 440:228–232.

Martinon, F., K. Burns, and J. Tschopp. 2002. The inflammasome: a molecular platform triggering activation of inflammatory caspases and processing of proIL-beta. Mol Cell. 10:417–426.

Mawuenyega, K.G., W. Sigurdson, V. Ovod, L. Munsell, T. Kasten, J.C. Morris, K.E. Yarasheski, and R.J. Bateman. 2010. Decreased clearance of CNS beta-amyloid in Alzheimer’s disease. Science. 330:1774.

McIntosh, T.K., D.H. Smith, D.F. Meaney, M.J. Kotapka, T.A. Gennarelli, and D.I. Graham. 1996. Neuropathological sequelae of traumatic brain injury: relationship to neurochemical and biomechanical mechanisms. Lab Invest. 74:315–342.

McKee, C.A., and J.R. Lukens. 2016. Emerging Roles for the Immune System in Traumatic Brain Injury. Front Immunol. 7:556.

Morganti-Kossmann, M.C., L. Satgunaseelan, N. Bye, and T. Kossmann. 2007. Modulation of immune response by head injury. Injury. 38:1392–1400.

Morimoto, K., J. Horio, H. Satoh, L. Sue, T. Beach, S. Arita, I. Tooyama, and Y. Konishi. 2011. Expression profiles of cytokines in the brains of Alzheimer’s disease (AD) patients compared to the brains of non-demented patients with and without increasing AD pathology. J Alzheimers Dis. 25:59–76.

Nicoll, J.A., R.E. Mrak, D.I. Graham, J. Stewart, G. Wilcock, S. MacGowan, M.M. Esiri, L.S. Murray, D. Dewar, S. Love, T. Moss, and W.S. Griffin. 2000. Association of interleukin-1 gene polymorphisms with Alzheimer’s disease. Ann Neurol. 47:365–368.

Nizamutdinov, D., and L.A. Shapiro. 2017. Overview of Traumatic Brain Injury: An Immunological Context. Brain Sci. 7.

O’Brien, W.T., L. Pham, G.F. Symons, M. Monif, S.R. Shultz, and S.J. McDonald. 2020. The NLRP3 inflammasome in traumatic brain injury: potential as a biomarker and therapeutic target. J Neuroinflammation. 17:104.

Parachikova, A., M.G. Agadjanyan, D.H. Cribbs, M. Blurton-Jones, V. Perreau, J. Rogers, T.G. Beach, and C.W. Cotman. 2007. Inflammatory changes parallel the early stages of Alzheimer disease. Neurobiol Aging. 28:1821–1833.

Pelegrin, P., and A. Surprenant. 2007. Pannexin-1 couples to maitotoxin- and nigericin-induced interleukin-1beta release through a dye uptake-independent pathway. J Biol Chem. 282:2386–2394.

Pimplikar, S.W. 2014. Neuroinflammation in Alzheimer’s disease: from pathogenesis to a therapeutic target. J Clin Immunol. 34 Suppl 1:S64–69.

Prokop, S., K.R. Miller, and F.L. Heppner. 2013. Microglia actions in Alzheimer’s disease. Acta Neuropathol. 126:461–477.

Roberts, G.W., S.M. Gentleman, A. Lynch, L. Murray, M. Landon, and D.I. Graham. 1994. Beta amyloid protein deposition in the brain after severe head injury: implications for the pathogenesis of Alzheimer’s disease. J Neurol Neurosurg Psychiatry. 57:419–425.

Rojo, L.E., J.A. Fernandez, A.A. Maccioni, J.M. Jimenez, and R.B. Maccioni. 2008. Neuroinflammation: implications for the pathogenesis and molecular diagnosis of Alzheimer’s disease. Arch Med Res. 39:1–16.

Royal, T., P. Ahluwalia, M. Gulhane, E.L. Salles, P. Gaur, M. Ahluwalia, T. Randhawa, S. Sunil, S. Budim, K. Akter, M.B. Khan, S. Ghosh, B. Baban, D.C. Hess, F.L. Vale, K.M. Dhandapani, R. Kolhe, and K. Vaibhav. 2026. Neuroinflammatory and functional outcomes after TBI are sex-dependent: Lessons from estrous-phase stratified female mice. Neurochem Int. 195:106136.

Saresella, M., F. La Rosa, F. Piancone, M. Zoppis, I. Marventano, E. Calabrese, V. Rainone, R. Nemni, R. Mancuso, and M. Clerici. 2016. The NLRP3 and NLRP1 inflammasomes are activated in Alzheimer’s disease. Mol Neurodegener. 11:23.

Schaffert, J., C. LoBue, C.L. White, H.S. Chiang, N. Didehbani, L. Lacritz, H. Rossetti, M. Dieppa, J. Hart, and C.M. Cullum. 2018. Traumatic brain injury history is associated with an earlier age of dementia onset in autopsy-confirmed Alzheimer’s disease. Neuropsychology. 32:410–416.

Schroder, K., R. Zhou, and J. Tschopp. 2010. The NLRP3 inflammasome: a sensor for metabolic danger? Science. 327:296–300.

Shaheen, M.J., A.M. Bekdash, H.A. Itani, and J.M. Borjac. 2021. Saffron extract attenuates neuroinflammation in rmTBI mouse model by suppressing NLRP3 inflammasome activation via SIRT1. PLoS One. 16:e0257211.

Smith, C., S.M. Gentleman, P.D. Leclercq, L.S. Murray, W.S. Griffin, D.I. Graham, and J.A. Nicoll. 2013. The neuroinflammatory response in humans after traumatic brain injury. Neuropathol Appl Neurobiol. 39:654–666.

Sutton, C.E., S.J. Lalor, C.M. Sweeney, C.F. Brereton, E.C. Lavelle, and K.H. Mills. 2009. Interleukin-1 and IL-23 induce innate IL-17 production from gammadelta T cells, amplifying Th17 responses and autoimmunity. Immunity. 31:331–341.

Sy, M., M. Kitazawa, R. Medeiros, L. Whitman, D. Cheng, T.E. Lane, and F.M. Laferla. 2011. Inflammation induced by infection potentiates tau pathological features in transgenic mice. Am J Pathol. 178:2811–2822.

Tan, M.S., J.T. Yu, T. Jiang, X.C. Zhu, and L. Tan. 2013. The NLRP3 inflammasome in Alzheimer’s disease. Mol Neurobiol. 48:875–882.

Thakkar, R., R. Wang, G. Sareddy, J. Wang, D. Thiruvaiyaru, R. Vadlamudi, Q. Zhang, and D. Brann. 2016. NLRP3 Inflammasome Activation in the Brain after Global Cerebral Ischemia and Regulation by 17beta-Estradiol. Oxid Med Cell Longev. 2016:8309031.

Villapol, S., D.J. Loane, and M.P. Burns. 2017. Sexual dimorphism in the inflammatory response to traumatic brain injury. Glia. 65:1423–1438.

Wallisch, J.S., D.W. Simon, H. Bayir, M.J. Bell, P.M. Kochanek, and R.S.B. Clark. 2017. Cerebrospinal Fluid NLRP3 is Increased After Severe Traumatic Brain Injury in Infants and Children. Neurocrit Care. 27:44–50.

Walsh, J.G., D.A. Muruve, and C. Power. 2014. Inflammasomes in the CNS. Nat Rev Neurosci. 15:84–97.

Webster, S.J., L.J. Van Eldik, D.M. Watterson, and A.D. Bachstetter. 2015. Closed head injury in an age-related Alzheimer mouse model leads to an altered neuroinflammatory response and persistent cognitive impairment. J Neurosci. 35:6554–6569.

Wo, J., F. Zhang, Z. Li, C. Sun, W. Zhang, and G. Sun. 2020. The Role of Gamma-Delta T Cells in Diseases of the Central Nervous System. Front Immunol. 11:580304.

Xu, X., D. Yin, H. Ren, W. Gao, F. Li, D. Sun, Y. Wu, S. Zhou, L. Lyu, M. Yang, J. Xiong, L. Han, R. Jiang, and J. Zhang. 2018. Selective NLRP3 inflammasome inhibitor reduces neuroinflammation and improves long-term neurological outcomes in a murine model of traumatic brain injury. Neurobiol Dis. 117:15–27.

Yi, H.J., J.E. Lee, D.H. Lee, Y.I. Kim, C.B. Cho, I.S. Kim, J.H. Sung, and S.H. Yang. 2020. The role of NLRP3 in traumatic brain injury and its regulation by pioglitazone. J Neurosurg. 133:1083–1091.

Youm, Y.H., R.W. Grant, L.R. McCabe, D.C. Albarado, K.Y. Nguyen, A. Ravussin, P. Pistell, S. Newman, R. Carter, A. Laque, H. Munzberg, C.J. Rosen, D.K. Ingram, J.M. Salbaum, and V.D. Dixit. 2013. Canonical Nlrp3 inflammasome links systemic low-grade inflammation to functional decline in aging. Cell Metab. 18:519–532.

Yu, X., W. Liu, S. Chen, X. Cheng, P.A. Paez, T. Sun, F. Yuan, C. Wei, J.W. Landry, A.S. Poklepovic, H.D. Bear, J.R. Subjeck, E. Repasky, C. Guo, and X.Y. Wang. 2021. Immunologically programming the tumor microenvironment induces the pattern recognition receptor NLRC4-dependent antitumor immunity. J Immunother Cancer. 9.

Zarruk, J.G., A.D. Greenhalgh, and S. David. 2018. Microglia and macrophages differ in their inflammatory profile after permanent brain ischemia. Exp Neurol. 301:120–132.

Zhang, Z., H. Wang, B. Tao, X. Shi, G. Chen, H. Ma, R. Peng, and J. Zhang. 2025. Attenuation of Blood-Brain Barrier Disruption in Traumatic Brain Injury via Inhibition of NKCC1 Cotransporter: Insights into the NF-kappaB/NLRP3 Signaling Pathway. J Neurotrauma. 42:814–831.

Zhu, S.G., J.G. Sheng, R.A. Jones, M.M. Brewer, X.Q. Zhou, R.E. Mrak, and W.S. Griffin. 1999. Increased interleukin-1beta converting enzyme expression and activity in Alzheimer disease. J Neuropathol Exp Neurol. 58:582–587.

Ziebell, J.M., and M.C. Morganti-Kossmann. 2010. Involvement of pro- and anti-inflammatory cytokines and chemokines in the pathophysiology of traumatic brain injury. Neurotherapeutics. 7:22–30.

